# Kmerator Suite: design of specific k-mer signatures and automatic metadata discovery in large RNA-Seq datasets

**DOI:** 10.1101/2021.05.20.444982

**Authors:** Sébastien Riquier, Chloé Bessiere, Benoit Guibert, Anne-Laure Bouge, Anthony Boureux, Florence Ruffle, Jérôme Audoux, Nicolas Gilbert, Haoliang Xue, Daniel Gautheret, Thérèse Commes

**Affiliations:** IRMB, University of Montpellier, INSERM, 80 rue Augustin Fliche, Montpellier, France; SeqOne, Montpellier, France; Institute for Integrative Biology of the Cell, CEA, CNRS, Université Paris Saclay, Gif sur Yvette, France

## Abstract

The huge body of publicly available RNA-seq libraries is a treasure of functional information allowing to quantify the expression of known or novel transcripts in tissues. However, transcript quantification commonly relies on alignment methods requiring a lot of computational resources and processing time, which does not scale easily to large datasets. K-mer decomposition constitutes a new way to process RNA-seq data for the identification of transcriptional signatures, as k-mers can be used to quantify accurately gene expression in a less resource-consuming way. We present the Kmerator Suite, a set of three tools designed to extract specific k-mer signatures, quantify these k-mers into RNA-seq datasets and quickly visualize large datasets characteristics. The core tool, Kmerator, produces specific k-mers for 97% of human genes, enabling the measure of gene expression with high accuracy in simulated datasets. KmerExploR, a direct application of Kmerator, uses a set of predictor genes specific k-mers to infer metadata including library protocol, sample features or contaminations from RNA-seq datasets. KmerExploR results are visualised through a user-friendly interface. Moreover, we demonstrate that the Kmerator Suite can be used for advanced queries targeting known or new biomarkers such as mutations, gene fusions or long non coding-RNAs for human health applications.

## INTRODUCTION

Publicly available human RNA-sequencing (RNA-seq) datasets are precious resources for biomedical research. RNA-seq data is widely used to identify actively transcribed genes, quantify gene or transcript expression, identify new fusion transcripts or identify alternative splicing or mutation events. The search for specific transcriptional events or RNAs across large-scale data has become essential in precision medicine. An increasing number of studies attempt to analyze in a retrospective fashion the vast repository of RNA-seq data, including normal and pathological conditions, to discover or validate RNA biomarkers for disease diagnosis (1, 2).

For this purpose, it is important to select relevant RNA-seq datasets with homogeneous characteristics and sufficient samples among thousands of publically available files. The reanalysis of RNA-seq datasets poses two major challenges. A first challenge is to filter data series and select the most homogeneous and reliable set of libraries for exploration in a context of incomplete metadata. A second challenge is to perform RNA biomarker quantification in reasonable time and with reasonable precision in such datasets. Alignment-based methods require significant computational resources, making them inadequate for querying datasets in the order of 100 or 1000 files for a specific biomarker. Pseudo-alignment algorithms like Kallisto (3) and Salmon (4) are much faster but depend on a complete transcriptome. This highlights the need for tools enabling fast and specific quantification of candidate sequences in a large set of RNA-seq data. Recently, approaches relying on k-mer from raw sequences files have emerged and are used for the query of transcriptomic data. These methods require less time and computational resources than common ones and are suited to various biological questions, including the analysis of unannotated and atypical RNA transcriptional events. For instance Okamura et al. proposed an ultrafast mRNA quantification method, based on unique k-mers, that outperforms conventional approaches (5). Yu et al. (6) investigated gene-fusion queries of all tumor samples from the TCGA project using k-mer sets. The DE-kupl pipeline developed by Audoux et al. (7) finds differential events between 2 groups of RNA-seq data at the k-mer level.

Although any transcript sequence can be decomposed into k-mers, only a subset of these k-mers is specific for the transcript. We call this subset the k-mer signature. These specific k-mers can then be quantified in RNA-seq raw data, making it quick and easy to measure the candidate transcript expression level in a wide range of RNA-seq datasets.

In this paper, we present the Kmerator Suite, a set of three tools designed to i/ extract k-mer signatures from transcripts, ii/ quantify these k-mers into RNA-seq datasets and iii/ visualize large RNA-seq datasets characteristics using precomputed signatures. The core of this suite is Kmerator, that generates k-mer signatures specific for genes or transcripts. A second tool, countTags, is used to quantify selected k-mers across raw RNA-seq files. We first tested the performance of Kmerator+countTags over the whole transcriptome and showed that k-mer signatures quantification results were close to simulated counts data. The third tool, KmerExploR, demonstrates the capacity of Kmerator+countTags pipeline combined to a set of predefined k-mer signatures, to perform metadata extraction from raw RNA-seq data. KmerExploR extracts sample characteristics related to the sequencing protocol (ribosomal depletion, polyA+, strand-specific protocol, 5’/3’ bias…), tissue origin (gender) as well as possible contaminations (mycoplasma, virus, other species or cell lines). Such high-level quality control procedures are valuable as a screening tool before analyzing datasets of uncertain quality such as public datasets. KmerExploR can also be used in advanced applications to look for user-defined transcripts resulting from mutated alleles or gene fusions in RNA-seq datasets.

## MATERIALS AND METHODS

### Kmerator: k-mer signatures identification

An overview of the Kmerator Suite is provided in Figure 1a. Kmerator is a tool designed for the prediction of specific k-mers from input sequences, considering a reference genome and an ENSEMBL-like fasta transcriptome (see Figure 1a and S1a). It is implemented in Julia programming language (https://julialang.org) and distributed with GitHub (https://github.com/Transipedia/kmerator). Kmerator strictly depends on a reference genome (fasta or Jellyfish (8) index format) and on an Ensembl fasta format transcriptome, to define a k-mer as specific or not, depending on the number of occurrences on each reference. The reference genome and transcriptome fasta, used in this paper, have been downloaded here: https://www.ensembl.org/info/data/ftp/index.html. The procedure also needs a list of gene/transcript Ensembl IDs (or gene symbols) or sequences in fasta format from which Kmerator will extract specific k-mers. As shown in figure S1a, Kmerator first uses the Jellyfish software to index and count k-mers from the reference genome and transcriptome. For both genome and transcriptome fasta files, Jellyfish produces a hash table including all possible k-mers and their number of occurrences. These hash tables are stored for further querying. Second, using Jellyfish query, Kmerator generates, for each input gene/transcript, the list of k-mers derived from this sequence and their corresponding genome and transcriptome counts. These k-mers are then filtered according to the following criteria: i/ only k-mers associated to a biological event (transcript or gene, splice-variant, chimeric RNA, circRNA, etc) are retained and ii/ k-mers must be specific according to Kmerator rules (see Figure 1c and S1a). Indeed, Kmerator includes 3 different levels of specificity (–level option), ‘gene’, ‘transcript’ and ‘chimera’, detailed below:

- gene level specific k-mers are found zero (to include k-mers containing splicing junctions) or one time in the reference genome. They are also present in the reference transcriptome in at least one isoform transcript sequence. If we want to select only k-mers matching at least *n* isoforms on a total of *N*, a threshold can be set to the proportion of isoforms n/N the k-mer has to be specific to, using the –threshold option.
- transcript level specific k-mers are found zero or one time in the reference genome. They also match the reference transcriptome only once (transcript specificity). If the candidate transcript is not annotated, the –unannotated option must be added. In this case, k-mers found zero or one time in the reference genome and that do not map to the reference transcriptome are retained.
- chimera level specific k-mers are neither found in the reference genome nor in the reference transcriptome. This level must be combined to the –unannotated option. Kmerator outputs the list of specific k-mers (also called k-mer gene/transcript signature) according to the chosen parameters in fasta format, for each input sequence.

**Figure 1.**
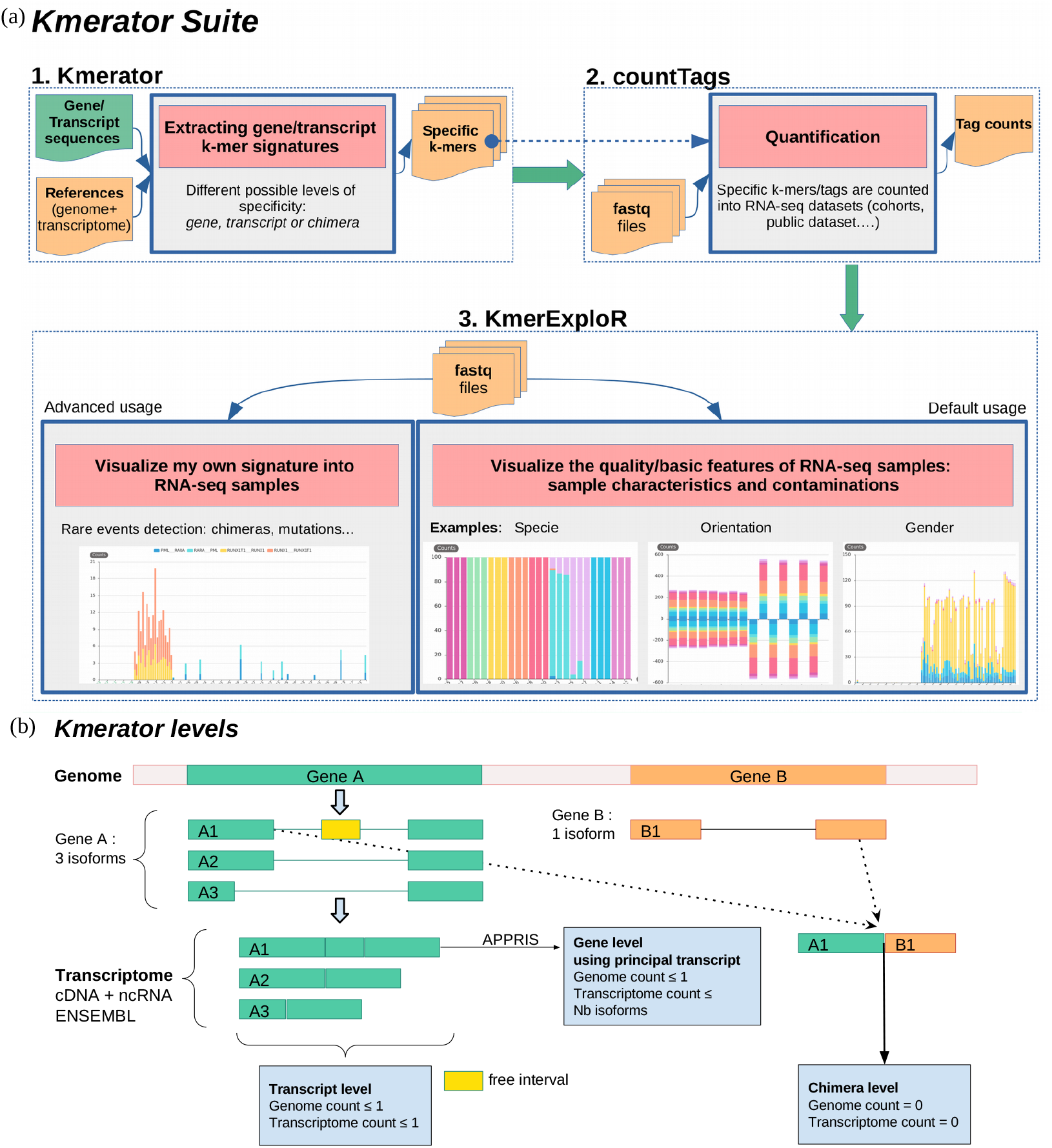
Kmerator Suite and Kmerator levels definition. (a) The Kmerator Suite is a set of 3 tools: 1. Kmerator extracts gene/transcript k-mer signatures. It takes as input a reference genome and a reference transcriptome + a list of gene or transcript sequences to extract specific k-mers from. The output is a set of fasta files (1 per input gene/transcript sequence) with the specific k-mers. 2. countTags quantifies input k-mers in a set of input sequencing raw files (fastq files) and outputs a count table. 3. KmerExploR is a particular application of Kmerator/countTags to visualize input RNA-seq datasets (set of fastq files) characteristics. The default usage includes characteristics related to the sequencing protocol (ribosomal depletion, polyA+, strand-specific protocol, 5’/3’ bias), tissue origin (gender) and possible contaminations (mycoplasma, virus, other species or HeLa cell line). Users can also visualize their own signatures with the advanced usage. Details are given in the manuscript and Figure S1. (b) Kmerator extracts gene/transcript k-mer signatures with 3 possible levels of stringency. This figure describes how the different levels are defined (transcript, gene or chimera) for 2 example genes A and B. Example gene A has 3 isoforms: A1, A2 and A3. A1 is the only one with a free interval, i.e. a region not covered by other isoforms, and is defined as the principal transcript (APPRIS database). Therefore, at the transcript level, each transcript has its own specific k-mers set, depending on its coverage with other isoforms. At the gene level, the principal transcript defined with the APPRIS database is used, and specific k-mers can be common to several isoforms. At the chimera level (example of A1-B1 fusion) the k-mer is not described into annotations.

#### Options

The k-mer length can be set using the –length option. In the present study, we used the default 31 nt k-mer length according to the literature (7). The level of specificity is chosen among ‘gene’, ‘transcript’, ‘chimera’ with the –level option. When using the gene level, APPRIS database (http://appris.bioinfo.cnio.es) can be queried to identify the ‘PRINCIPAL’ transcript, using the –appris option. APPRIS defines as the ‘PRINCIPAL’ isoform a CDS (coding sequence) variant for each gene, based on the range of protein features. When this option is not used or no PRINCIPAL sequence is given by APPRIS (i.e. for lncRNA), the isoform with the longest sequence is kept. In this study, we always used the gene level in combination with the –appris option.

#### Kmerator usage on the entire transcriptome for performance assessment

Kmerator was tested to extract k-mer signatures from the whole human ENSEMBL transcriptome (combination of cDNA and ncRNA fasta files, version 91). The ENSEMBL transcriptome reference was filtered to remove any transcript with alternate loci (labels by “_alt”) and have been processed by Kmerator at both transcript (i.e. 199,181 transcripts) and gene (54,874 genes) levels with the –appris option previously described. At the transcript level, 62 transcripts have been ignored due to their length inferior to the k-mer length (31 nt). The processing on the whole transcriptome has been completed in less than 3 days at the gene level and 24h at the transcript level, using a LINUX server with 30 computing cores and 20 GB hard disk space.

### K-mer counting and expression quantification

#### Simulated data

To test the precision of k-mers quantification, we created a set of 10 simulated RNA-seq data. We first used the R *compcodeR* package (9) and “generateSyntheticData” function to simulate a count matrix with two conditions with 5 samples in each (samples.per.cond = 5). Each line of this matrix corresponds to a transcript of the Ensembl v91 annotation. Counts of transcripts with a length equal or inferior to 200 nt were not simulated. To highlight the quantification process, we increased the number of differentially expressed genes (n.diffexp = 10,000) with balanced over- and under-expressed fractions (fraction.upregulated = 0.5) and with authorised different dispersions between the conditions (between.group.diffdisp = TRUE, fraction.non.overdispersed = 0). Besides, we set the sequencing depth by RNA-seq file to 100M reads (seq depth = 100000000) and we didn’t filter low counts (filter.threshold.total = 0). Providing this dataframe and the Ensembl reference transcriptome, we used “simulate_experiment_countmat” function, from *polyester* R package (10), to generate paired-end and strand-specific (fr fashion) RNA-seq reads in fasta format. Finally, the fasta files have been converted to fastq.gz format using seqtk (https://github.com/lh3/seqtk).

#### CountTags

K-mers designed by Kmerator on the whole transcriptome were counted into the 10 simulated RNA-seq data. For this purpose, the list of k-mers was submitted to countTags (https://github.com/Transipedia/countTags), a tool written in C language (see Figure 1a). CountTags searches for short sequences (< 32nt) and their reverse complement with an exact match in fastq files and counts their occurences. We used a k-mer length of 31 nt (-k 31) and the paired-end option (–paired). Counts were normalized per billion of k-mers present in the dataset using the –kbp option.

#### Comparison with Kallisto

We compared the Kmerator + countTags pipeline to Kallisto regarding the performances in transcript/gene expression quantification on simulated data detailed above. As our pipeline can not quantify genes/transcripts without specific k-mers, we limited Kallisto quantification to the genes/transcripts having specific k-mers. Kallisto 0.43.1 (3) was run using the –fr-stranded option with the Ensembl 91 annotation file. For each pipeline, TPM (Transcripts Per Million) counts were compared to true normalized TPM using the Spearman’s correlation, either at the transcript or gene level. Counts estimated by Kallisto were merged at gene-level by summing normalized transcript counts.

### KmerExploR: exploring large RNA-seq datasets

KmerExploR is a command line tool powered by the back-end pipeline Kmerator + countTags. KmerExploR provides k-mers quantification results in RNA-seq samples as a graphical and user-friendly html interface (see Figure 1a). To deal with data heterogeneity and the weaknesses of RNA-seq technology, we developed a turnkey application using KmerExploR. Characterisation of a requested RNA-seq dataset can be improved with the quantification of selected genes (predictor genes) via Kmerator + countTags pipeline. Predictor genes and their corresponding specific k-mers are included into KmerExploR and have been selected based on literature to answer specific biological questions:

- Are my RNA-seq data based on poly-A selection protocol or ribo-depletion?
- Are my RNA-seq libraries stranded or not?
- What is/are the gender(s) corresponding to my samples?
- Is there a read coverage bias from 5’ to 3’ end along my dataset transcripts?
- Are my RNA-seq data contaminated by HeLa (presence of HeLa-derived human papillomavirus 18), mycoplasmas or other viruses such as hepatitis B virus?
- What is/are the species present into my samples?

#### Implementation

KmerExploR is a command line tool written in python 3. It can be installed on a server or on a personal computer from github or with pip command (see https://github.com/Transipedia/kmerexplor). No additional modules are required. KmerExploR does not need a lot of memory and can be launched from a laptop. It includes countTags, described above. From input fastq files, KmerExploR run countTags, with a multithreading option, to quantify built-in k-mers selection associated to each predictor gene. The detailed diagram is shown in Figure S1b. KmerExploR can also directly take countTags output files, as for large datasets, it could be useful to separately run countTags on a cluster for example. KmerExploR outputs an HTML file with css and javascript in separate files, using the echartsjs library to display user-friendly information (https://echarts.apache.org/en/index.html). Categories to show are described either in the built-in config file or in the user personal config file. KmerExploR also produces a tabulated text file with mean counts for each predictor gene in each category (rows) and in each sample (columns).

#### Predictor genes selection

We selected a subset of housekeeping genes from the list previously published by Eisenberg and Levanon (11) as well as some widely expressed histone genes which produce non-polyadenylated transcripts barely detected into poly-A+ RNA-seq (see Table 1). We also selected specific genes from chromosome Y that have an ubiquitous expression, from Maan’s et al. publication (12). For these different sets of genes, we designed specific k-mers using Kmerator at the gene level and also computed the k-mers reversed complementary counterparts for the orientation category. Housekeeping genes ubiquitous expression profile in various tissues, chromosome Y genes specific expression pattern in male tissues and histone genes low expression in polyA+ RNA-seq samples have been validated by exploring the GTEx database (https://www.gtexportal.org) (see Figure S2).

**Table 1.** List of predictor genes, by category, included in KmerExploR and associated RNA-seq dataset names used in this paper. The samples included in each dataset and some metadata are detailed in the supplementary table S1.

|  | Datasets | Predictor genes | Total k-mers number | References and details |
| --- | --- | --- | --- | --- |
| **PolyA/RiboD** | Dataset-FEATURES | HIST2H2AC, HIST2H2AB, HIST1H4J, HIST1H4I, HIST1H4F, HIST1H4D, HIST1H4C, HIST1H4B, HIST1H3I, HIST1H3H, HIST1H3G, HIST1H3F, HIST1H3E, HIST1H3C, HIST1H3B, HIST1H3A, HIST1H2BN, HIST1H2BM, HIST1H2BL, HIST1H2BH, HIST1H2BF, HIST1H2BE, HIST1H2BB, HIST1H2BA, HIST1H2AK, HIST1H2AH, HIST1H2AG, HIST1H2AB, HIST1H1T, HIST1H1E, HIST1H1D, HIST1H1B, HIST1H1A | 24512 | Figure S2 |
| **Orientation** | Dataset-FEATURES | VPS29, SNRPD3, REEP5, RAB7A, PSMB4, PSMB2, GPI, EMC7, CHMP2A, C1orf43, VPS29_rev, SNRPD3_rev, REEP5_rev, RAB7A_rev, PSMB4_rev, PSMB2_rev, GPI_rev, EMC7_rev, CHMP2A_rev, C1orf43_rev | 36638 | Figure S2<br>Eisenberg et al. (2013) |
| **Gender** | Dataset-FEATURES | UTY, TMSB4Y, TBL1Y, RPS4Y1, NLGN4Y, EIF1AY, DDX3Y | 21996 | Figure S2<br>Maan et al. (2017) |
| **5'/3' bias** | Dataset-FEATURES | VPS29, SNRPD3, REEP5, RAB7A, PSMB4, PSMB2, GPI, EMC7, CHMP2A, C1orf43 | 12705 | Figure S2<br>Eisenberg et al. (2013) |
| **Mycoplasma** | Dataset-MYCO | Mycoplasma_orale, Mycoplasma_hyorhinis, Acholeplasma_laidlawii, Mycoplasma_hominis, Mycoplasma_argininis, Mycoplasma fermentans | 363025 | Drexler et al. (2002)<br>SILVA Database |
| **Virus** | Dataset-VIRUS-CCLE | Human_gammaherpesvirus_4, Human_herpesvirus_4, Human_herpesvirus_8, Murine_leukemia_virus, Hepatitis_C_virus_genotype, Human_immunodeficiency_virus_1, Human_T_lymphotropic_virus_1, Squirrel_monkey_retrovirus, Human_T_lymphotropic_virus_2, Human_papillomavirus_type_92, Hepatitis_B_virus_strain, Human_immunodeficiency_virus_2, MuLV_related_virus_22Rv1/CWR, Bovine_viral_diarrhea_virus | 516882 | Uphoff et al. (2019) |
| **HeLa** | Dataset-HELA-CCLE | L1_mut7486, L1_mut7258, L1_mut6842, L1_mut6625, L1_mut6460, L1_mut6401, L1_mut5875, E7_mut806, E7_mut751, E6_mut549, E6_mut485, E6_mut287, E6_mut104, E1_mut2269, E1_mut1994, E1_mut1843, E1_mut1807, E1_mut1353, E1_mut1012 | 589 | Cantalupo et al. (2015) |
| **Species** | Dataset-SPECIES | Homo_sapiens_MT_CO1, Danio_rerio_mt_co1, Zea_mays_COX1, Saccharomyces_cerevisiae_COX1, Rattus_norvegicus_Mt_co1, Mus_musculus_mt_Co1, Gallus_gallus_MT_CO1, Drosophila_melanogaster_mt_Co1, Caenorabditis_elegans_ctc_3_MTCE, arabidopsis_thaliana_COX1 | 12119 | MT-CO1 (and orthologs) |
| **Chimeras** | Dataset-LEUCEGENE | PML-RARA, RARA-PML, RUNX1T1-RUNX1, RUNX1-RUNX1T1 | 724 |  |
| **lncRNA** | Dataset-LEUCEGENE | NONE | 78 | F. Rufflé et al. (2017) |
| **mutations** | Dataset-LEUCEGENE | TET2, KRAS, CEBPA | 10 |  |

For the detection of 5’ 3’-end biases, we used the specific k-mers from ubiquitous genes (orientation set) and individually attributed them to their corresponding region, 5’ UTR, 3’ UTR or CDS, depending on their position in the principal transcript, according to APPRIS database. For that purpose, we used ensembl annotations with the biomaRt R package which gives the information of the UTRs and CDS regions for each transcript. We searched the k-mers in transcripts CDS and UTRs sequences to label them by region. For mycoplasma tags selection, we first selected the most frequent mycoplasma found in cell contamination according to Drexler et al. (13). We then downloaded ribosomal RNA (rRNA) sequences of the 6 selected mycoplasma species from SILVA database v132 (14), which provides updated and curated rRNA sequences from Bacteria, Archaea and Eukaryota. Some species have several associated strains and therefore several rRNA sequences. We have included them all for the k-mers design. For HeLa detection, we selected HPV-18 transcripts reported to be expressed in HeLa cells (15). Using UGENE software (16), we manually modified these transcripts to match the mutations reported as HeLa specific in Cantalupo et al. study (15). We then defined sequences taking 30 nt on both sides of each mutation, before passing them to Kmerator to keep only k-mers not present in the human genome and transcriptome. For species identification, we selected the ones principally found in the SRA database. We then downloaded MT-CO1 human gene sequence and his orthologs in each of the selected species, using the corresponding animal reference genome and transcriptome sequences (Ensembl v91 for each). Finally, sequences of virus genomes have been downloaded from RefSeq using the common virus list provided by Uphoff et al. (17). All these potential contamination sequences were used to produce specific k-mers using Kmerator at the chimera level, to select tags that can be found neither in the human reference genome nor in the transcriptome. For the advanced application of KmerExploR, we designed k-mers corresponding to new or rare transcriptional events detected in Leucegene dataset (https://leucegene.ca/). For chimera detection, we used two well-known fusion RNA examples associated to chromosomal translocation and their reciprocal counterparts (RUNX1-RUNXT1 t(x,21) RUNXT1-RUNX1, PML-RARA t(15, 17) and RARA-PML). Specific k-mers are designed with Kmerator on 60 bp sequences spanning the junction. For mutation detection, we manually designed 31 bp k-mers centered on the mutation for reference and alternative sequences of 3 genes currently used in AML diagnosis: TET2, KRAS and CEBPA. We finally designed k-mers with Kmerator at the transcript level for a new long non coding RNA previously published in Ruffle et al. (18) as NONE “chr2-p21”.

#### RNA-seq dataset

In this paper, we illustrated KmerExploR output on several datasets, depending on the biological question, all described in the supplementary Table S1. Characteristics related to RNA-seq protocol, which we call basic features, are tested on 103 paired-end samples from ENCODE (Dataset-FEATURES). For the contaminations part, we used the PRJNA153913 study previously described as highly contaminated by mycoplasma (Dataset-MYCO). We also selected 3 public RNA-seq samples by species to check the relevance of our species specific k-mers (Dataset-SPECIES). HeLa contamination was tested in 3 cervical cancer CCLE cell lines: 1 HeLa and 2 negative controls (Dataset-HELA-CLE). Finally, for virus detection we used a subset of 22 samples from the CCLE dataset reported by Uphoff et al. (17) as contaminated (Dataset-VIRUS-CCLE).

## RESULTS

### Kmerator performances

To assess the Kmerator methodology, we first extracted k-mer signatures from all the human ENSEMBL transcriptome (i.e. 199,181 transcripts) and genes (i.e. 54,874 coding and noncoding genes). We were able to identify specific k-mers (k=31 nt) for 83% of human transcripts and 97% of human genes as shown in Figure 2a-b.

**Figure 2.**
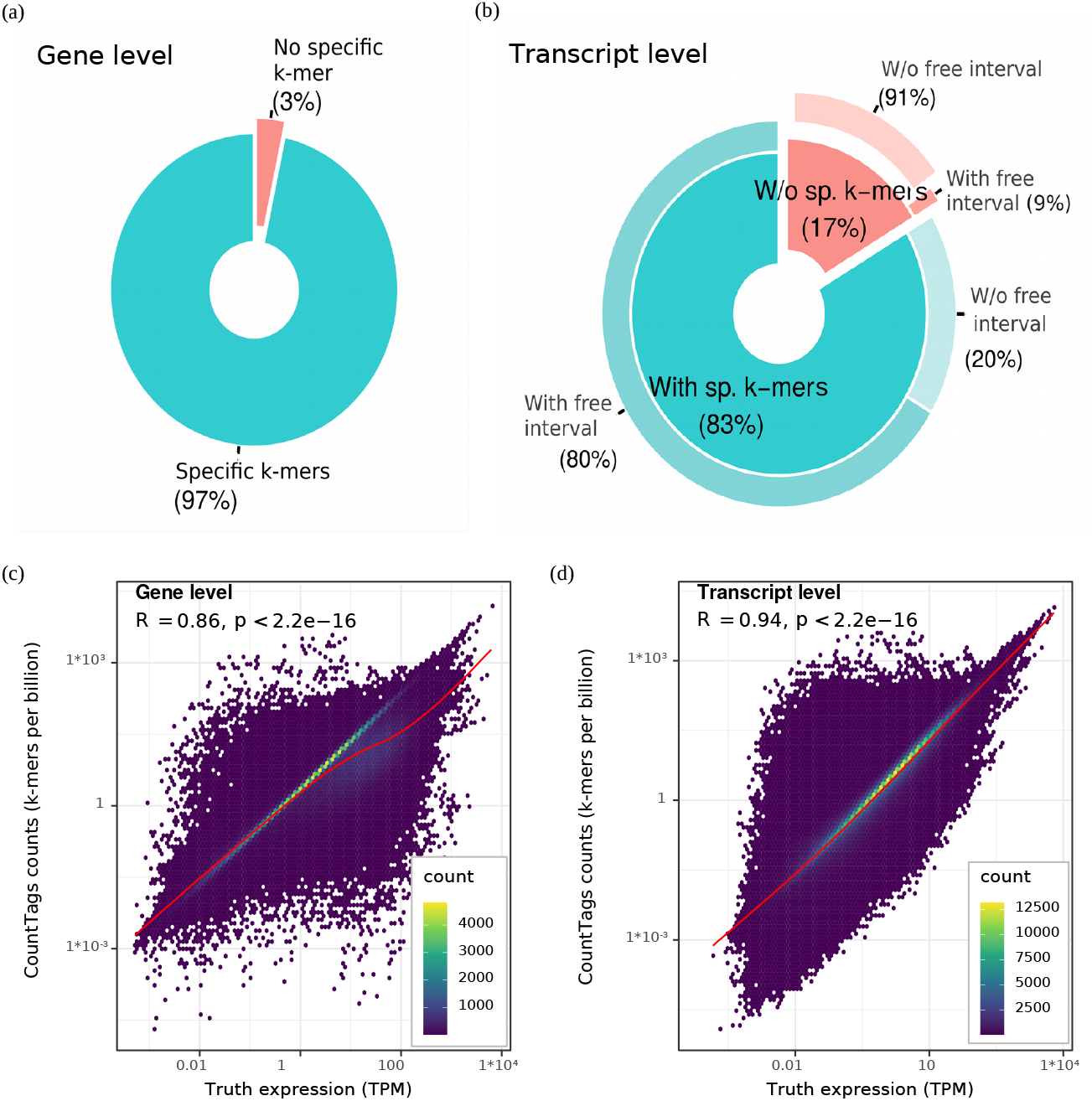
Kmerator performances on the whole transcriptome. We extracted k-mer signatures from all the human ENSEMBL transcriptome v91 at both gene (54,874 coding and non-coding genes, left) and transcript (i.e. 199,181 transcripts, right) levels. (a) The first pie chart represents the proportion of genes having specific k-mers (turquoise) versus the ones without specific k-mers (red). (b) In the same way, we represented the proportion of transcripts having specific k-mers (turquoise) or not (red). For these 2 classes, we looked at the percentage having free intervals, i.e regions in the transcript not shared with other isoforms (secondary pie). Most of the transcripts lacking specific k-mers don’t have free intervals (91%). (c,d) We tested Kmerator sensitivity to quantify simulated data, at both gene (c) and transcript (d) levels. We represented the k-mer counts normalized per billion of k-mers in the sample (Y axis) function of the true expression in TPM (X axis), on the whole simulated dataset. R is the spearman correlation coefficient between k-mer counts and TPM. Each point on the graph is a transcript and the color scale depends on the transcripts density on the graph.

This way, the transcriptome information has been almost entirely summarised by 69 760 957 k-mers at the transcript level and 88 003 855 at the gene level, corresponding to 23.8% and 30% of the total number of k-mers in the reference transcriptome respectively. The attribution of specific k-mers at the gene and transcript levels are fundamentally different: whereas the gene level (–appris option) accepts specific k-mers shared with other isoforms, the transcript level is more stringent and eliminates each k-mer shared by other ones. This explains the higher percentage of transcripts without specific k-mer compared to the gene level. To explain the absence of specific k-mers for some transcripts, we used BiomaRt genomic intervals to calculate the part of each transcript not covered by other isoforms, considering the strand, and named it “free interval” (see Figure 1b). As expected, 91% of transcripts without specific k-mer have no “free interval”, which means that they are completely covered by other transcripts, thus confirming the validation of the Kmerator process. The set of specific k-mers designed with Kmerator strongly depends on the input sequence and on the level of selection. At the gene level, we observed that the length of the input sequence was correlated with the number of designed specific k-mers (R = 0.91, *p*< 2.2e —16, see Figure S3a) but not at the transcript level (R =0.22, *p*< 2.2e —16, see Figure S3b). On the contrary, the transcript level depends on the overlap between the input transcript and the different isoforms. A high number of isoforms is correlated to a low number of specific k-mers (R = 0.79, *p*< 2.2e —16, see Figure S3c) and, in addition, the length of free intervals is strongly correlated to the number of specific k-mers (R = 0.94, *p*< 2.2e −16, see Figure S3d). Finally, k-mers design differs between biotypes and selection levels: the biotypes without specific k-mers mainly correspond to small RNAs (miRNAs, rRNA) at the gene level (see Figure S3e) and to coding and pseudo-genes at the transcript level (see Figure S3f).

Kmerator being designed for rapid large scale quantification in RNA-Seq data, we tested its accuracy to estimate gene and transcript expression using simulated data (see Materials and Methods). We have run Kmerator and countTags to search for all human genes and transcripts expression levels in a set of 10 simulated data. We assessed Spearman’s correlation between normalized k-mer counts and the ground truth. We used countTags k-mers mean count per transcript reported to the total of k-mers contained in the input fastq. As shown in Figure 2, the Spearman’s correlation factor comparing Kmerator+countTags results to the truth is 0.86 for the gene level (Figure 2c) and 0.94 for the transcript level (see Figure 2d), indicating a highly positive relationship with normalized counts (*p*<2*e*–16).

Quantification results are comparable when using the Kallisto pseudoalignment method, despite slightly higher correlation factors (gene and transcript R = 0.97, see Figure S4a-b). This result is consistent with the recent paper describing Matataki (5), another quantification tool based on k-mers. Our pipeline being not specifically dedicated to gene quantification but for rapid exploration of large datasets, it is accurate enough to evaluate gene and transcript expression levels in RNA-seq data. Interestingly, the precision of Kallisto quantification decreases strongly with transcripts/genes not covered by Kmerator (see Figure S4c-d), showing that each protocol using k-mer principle struggles to correctly quantify sequences that do not possess distinctive k-mers.

Finally, we tested speed performance of countTags processing time on random subparts of a sample simulated data (10M, 101nt paired-end reads), while increasing the number of quantified k-mers (1/1000/1M). It appears that processing time remains low compared to alignment based protocols (about 1min for 10M reads) and depends on the number of k-mers quantified (see Figure S4e). These results support an optimized usage of the Kmerator Suite protocol for its primary usage: the research of a limited number of signatures in large RNA-seq datasets.

### KmerExploR for inspecting large RNA-seq datasets

We developed KmerExploR to improve the characterisation of a large RNA-seq datasets using the quantification of selected predictor genes. Predictor genes have been selected based on literature to answer specific questions (see Table 1). As described in Materials and Methods, we first extracted with Kmerator, sets of specific k-mers from gene sequences and use KmerExploR to count the k-mer occurrences into RNA-seq datasets and visualise the results. Here, we present the results obtained with specific datasets (Table 1 and table S1) selected to highlight the rapid control of biological and technical parameters using KmerExploR.

The results of the basic features, including sample gender, polyA or ribo-depletion, orientation and 5’/3’ bias are presented in see Figure 3. As previously described, sample gender is determined by searching for k-mers corresponding to genes located on the Y chromosome. The k-mer signature clearly separates samples depending on the gender. We can however mention the Y chromosome genes expression dispersion between the samples, that can be explained by the variability of cell types and public RNA-seq experiment parameters, including sequencing depth and methods of RNA extraction and selection. For instance, the 4 male samples with the lowest expression (ENCFF232KGN, ENCFF434EMO, ENCFF831HCD and ENCFF992HBZ) comes from a unique study (ENCSR999ZCI).

**Figure 3.**
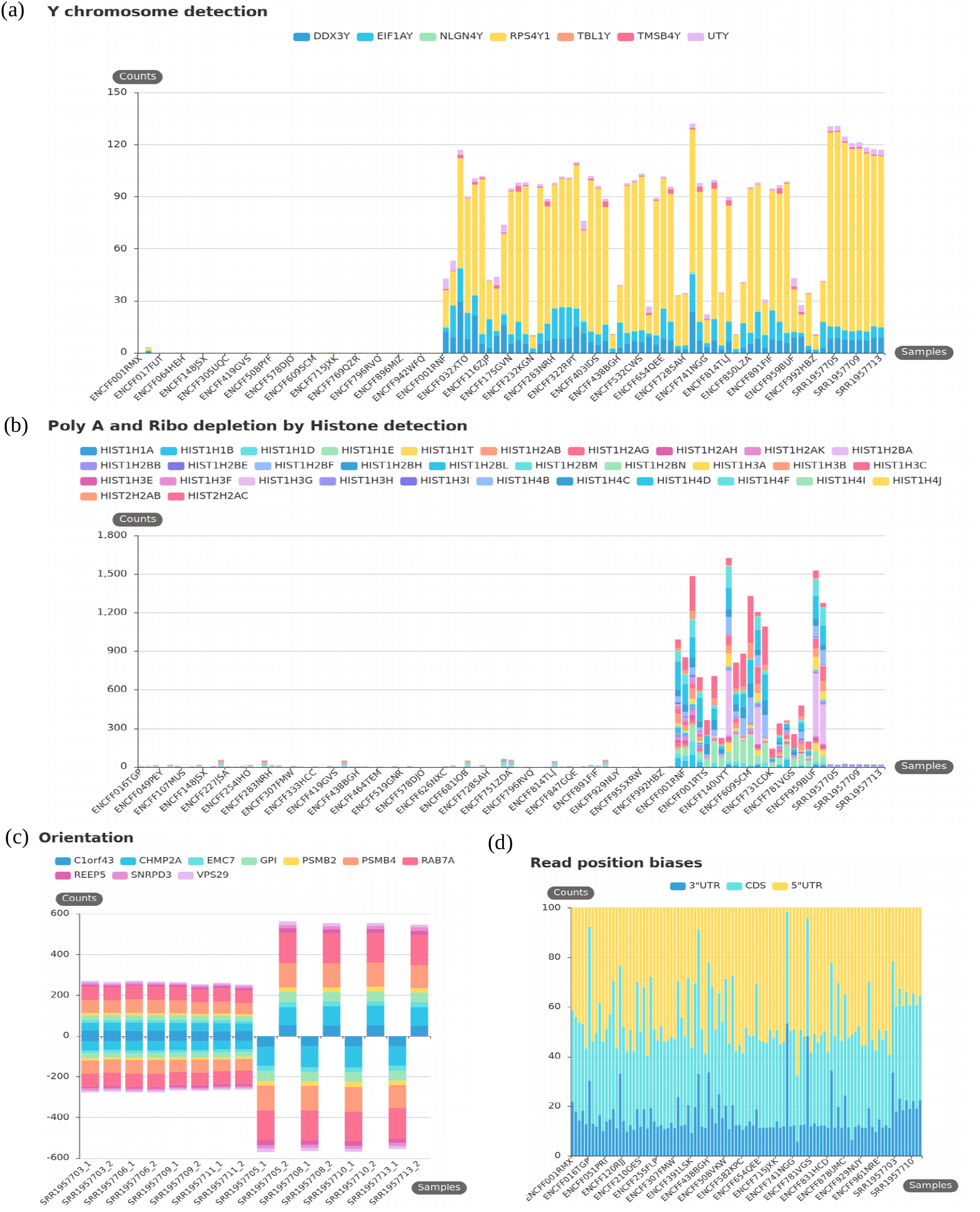
KmerExploR default usage: basic features. All barplots are generated from Dataset-FEATURES described in table S1 (103 paired-end ENCODE samples) except for the orientation (c) which is a subset of 8 RNA-seq from Dataset-FEATURES. For each barplot, the legend lists the set of predictor genes for which k-mers mean counts are computed (see also table 1). Samples are on the X axis. (a), (b) and (c) have the mean k-mer counts by gene normalized per billion of k-mers on the Y axis. (a) Sex determination. Samples are sorted by gender in the order female then male. (b) PolyA+ selection versus Ribo-depletion by histones detection. Samples are sorted by protocol in this order: polyA, ribo-depletion, unknown. (c) Stranded versus unstranded sequencing protocol. For this category, both fastq files by sample are shown. The 4 first samples are unstranded and the 4 last stranded. (d) Read position biases along 5’UTR, CDS and 3’UTR regions. After computing k-mers mean counts by gene, they are summed up by region 5’UTR, 3’UTR or CDS and converted in % (Y axis).

Gene abundance can be measured in RNA-seq data through sequencing of mRNA or ribo-depleted total RNA samples. The mRNA protocol relies on polyA selection, when the total RNA method is based on rRNA depletion (Ribozero protocol). Though, non-polyadenylated transcripts should only be found in data produced using this procedure, when they should barely be detectable in mRNA samples. As the majority of Histone transcripts are known to be non-polyadenylated, we used this characteristic first to detect sample contamination by non polyadenylated RNA, and secondly to infer from the result the RNA preparation procedure. We first investigated the expression level of all histone genes and retained the most highly expressed according to the literature. Secondly, we analysed their expression pattern using the GTEX resource. As RNA-seq from GTEX are exclusively produced from polyA selected RNAs samples, we used this database to select histone genes showing the lowest expression levels (see Figure S2b). We used this set of histone genes to test a selection of ENCODE samples which metadata indicates either polyA or ribo-depletion protocol (Supplementary table S2). The results clearly demonstrate differences between libraries prepared by ribo-depletion vs polyA selection for most of the chosen histone genes. We observe histone genes expression variability between the samples demonstrating again the disparity of public data. Strand-specific and unstranded library preparation are two commonly used preparation protocols that differ by their ability to retain or not RNA strand information. To detect this characteristic from RNA-Seq data, we designed kmers, specific for a set of ubiquitous genes (Table 1) and their reverse complement counterparts. K-mers on the forward strand are counted as positive and their reverse complement as negative, permitting to determine the orientation of the library. If forward and reverse tags are found in equivalent proportions in the same fastq file, data are considered as “unstranded”. This leads graphically to a balanced distribution between positive and negative counts. As shown in figure 3, using this property we are able to clearly separate unstranded and stranded libraries. 5’ to 3’-end bias is a difference of reads repartition along the transcripts, classically linked to library preparation: incomplete retro-transcription or specific protocols. A comparison between polyA selection and ribo-depletion protocols has previously shown coverage differences across transcripts with a poor 5’-end coverage with polyA selection method (19). Knowing whether an RNA-seq sample possesses a read repartition bias is critical for isoforms detection, or simply to give an indication on the library construction protocol used in large-scale analysis of public data. Using previously described housekeeping genes (Table 1), we have selected different sets of specific k-mers depending on their position into the regions defined as 5’ UTR, 3’ UTR and CDS. The Figure 3c shows the repartition in percent of these k-mers across the Dataset-FEATURES samples. Representing the mean k-mer counts as a percentage allows us to evaluate the distribution homogeneity across 5’, 3’ and CDS regions between the 103 ENCODE samples. This global representation grouping together several genes allows us to identify samples for which one region has a very little coverage. Here, 4 samples have less than 10% 5’UTR coverage (ENCFF734ZAD, ENCFF770NYA, ENCFF419GVS and ENCFF016TGP). we can also notice a better homogeneity of coverage for ribodepleted samples.

### Detection of potential contamination

Different microorganisms like mycoplasma and virus can contaminate samples and cell cultures, modifying the metabolism of the cell and therefore biases the results of ensuing analysis. Moreover, cancer research has shown that viruses are responsible for about 20% of human cancers (20). To detect contaminants in RNA-seq data, tools relying on alignment like DecontaMiner (21) or viGEN (22) have been widely used, but the alignment step is time and memory consuming. Exact alignment of k-mers based approaches like Kraken (23) and Taxonomer (24) are an alternative for taxonomic classification. However, these tools are complex and involve data cleaning from adaptators (trimming), the use of internal and external databases and/or probabilistic models for contaminant classification. Using a specific and reduced set of k-mers, we have seen an advantage to quickly detect principal contaminants of human cells in RNA-seq datasets, free from alignment methods.

Because mycoplasma is a common source of cell culture sample contamination and could affect host gene expression (25), we choose to control its presence in RNA-seq data. Mycoplasma contamination is evaluated through the detection of specific k-mers corresponding to 16S rRNA sequences according to the literature. In fact, in the Olarerin-George and Hogenesch study, they showed that 90% of the specific mycoplasma-mapped reads from human RNA-seq samples mapped to mycoplasma ribosomal RNA. We selected 6 species that have the highest record rate of detection in cell culture samples (i.e. A. laidlawii, M. fermentans, M. hominis, M. hyorhinis, M. orale and M. arginini) (13) to design our k-mers. We used an RNA-Seq data series previously described as highly contaminated (25) (study PRJNA153913) to test the relevance of our approach. As shown in Figure 4, we can easily detect the 6 selected mycoplasma species in some samples, with a prevalence for the M. hyorhinis species. Comparing our results with Olarerin-George and Hogenesch study which use Bowtie 1 alignment and BLAST+ to filter non-specific reads, we were able to confirm mycoplasma ribosomal RNA presence for the same samples (see Figure S5a). Moreover, we observe a high proportionality between our k-mer counts and their read counts on the 33 single-read samples (Dataset-MYCO described in table S1), for each of the 6 common Mycoplasma species.

**Figure 4.**
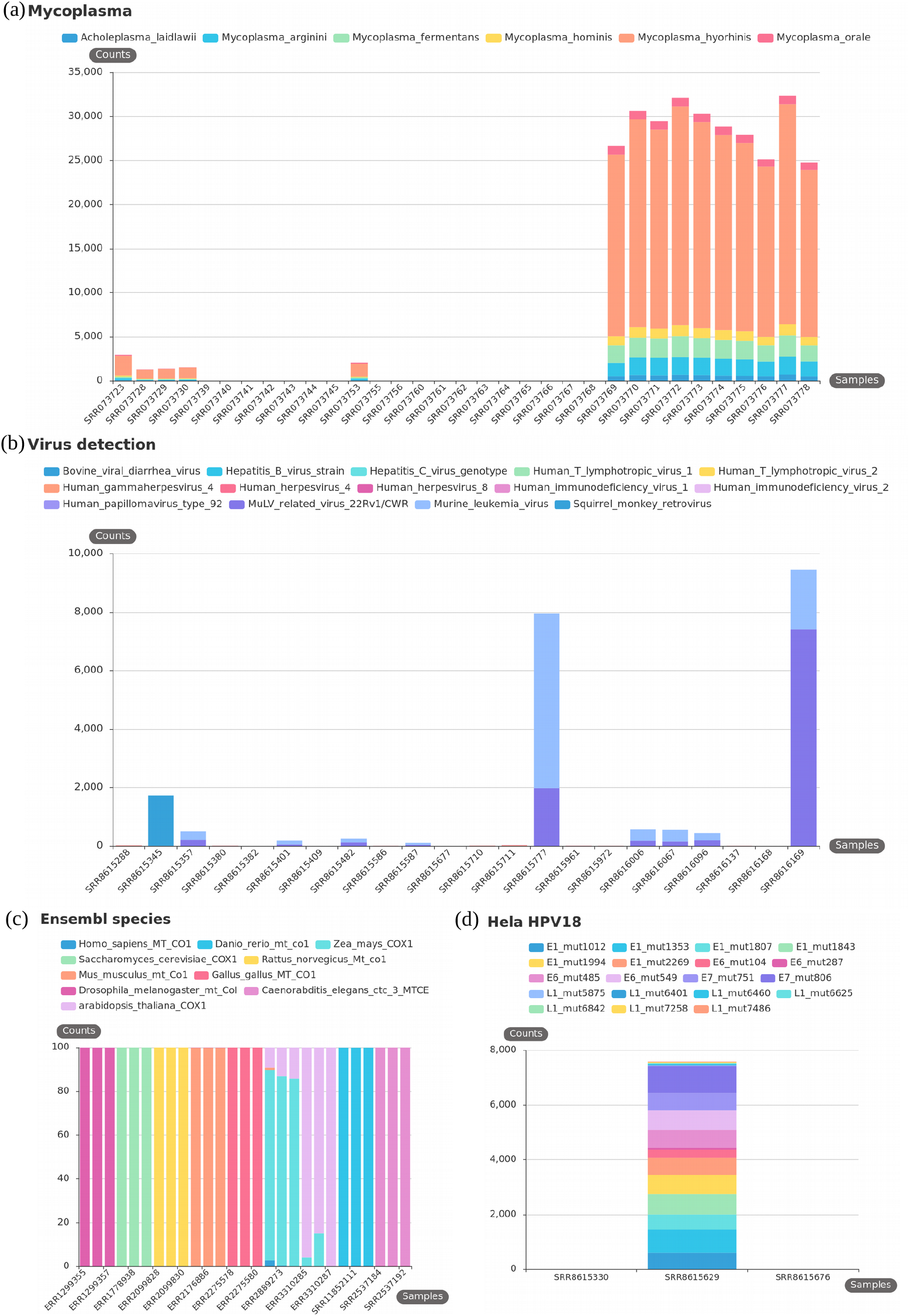
KmerExploR default usage: contaminations. All barplot datasets are described in table S1. For each barplot, the legend lists the set of predictors for which k-mers mean counts are computed (details in table 1). Samples are on the X axis. (a), (b) and (d) have the mean k-mer counts by gene normalized per billion of k-mers on the Y axis. (a) Mycoplasma contamination on the Dataset-MYCO (33 single-read samples). (b) Virus detection on the Dataset-VIRUS-CCLE (22 paired-end samples). (c) Species determination on the Dataset-SPECIES (27 paired-end samples). For this category, after computing k-mers mean counts by species, they are converted in % (Y axis) to avoid big expression differences between species. (d) HeLa determination on the Dataset-HELA-CCLE (3 paired-end samples). The sample in the middle is an HeLa cell line and the 2 others are negative controls (SF767 and siha cells).

Viruses are a significant cause of human cancers. Several studies interrogate for the presence of major viruses known to infect human and other mammalian cells (17, 26, 27). Recently, Uphoff et al. screened more than 300 Cancer Cell Line Encyclopedia RNA-seq data using the Taxonomer interactive tool and compared the results to virus specific PCR analysis, revealing 20 infected cell lines with different viruses (17). To rapidly explore the potential presence of viruses into RNA-seq datasets with our k-mer based approach, we used the same virus reference genomes as described in Uphoff et al. study. Using Kmerator at the chimera level (absent from human annotations), we designed specific k-mers for each virus and searched them in a subset of contaminated CCLE data according to Uphoff et al. and into negative controls, to validate our protocol ability to detect viruses. Results are shown in the Figure 4b and S5b. Our results are consistent with C. Uphoff et al. study using both PCR validation and dedicated detection tools.

HeLa is the first immortal human cell line, coming from Henrietta Lacks cancerous tissue samples. Her cancer was triggered by an infection with human papillomavirus type 18 (HPV-18). Nowaday, this cell line is largely used in medical research. Looking for several viruses into public RNA-seq cancer-related databases revealed the presence of HPV-18 sequences in many cancers (28) which closely resemble the HPV-18 viral sequence that is integrated into HeLa cells, suggesting a contamination. 3 segments of HPV-18 are integrated into HeLa genome on chromosome 8 and include the long control region (LCR), the E6, E7 and E1 genes, and partial coding regions for the E2 and L1 genes [16]. These genes are expressed in HeLa cells, and mutations have been found specifically in HeLa cells. Thus, selecting these mutated HeLa HPV-18 genes specific k-mers and counting them into 3 CCLE RNA-seq datasets (1 positive sample and 2 negative controls) we validated the accuracy of our selection as we are able to find our k-mers selection specifically in HeLa cells. We also checked the results in other HeLa samples from the PRJNA639358 study (see Figure S5e).

As for HeLa cells, cross-species contamination remains a documented “danger” for results interpretation in molecular biology (29). The probability of mixed cell lines in samples preparation, usage of polymerase chain reaction (PCR) which can accidentally amplify the wrong piece of DNA, plus an unknown probability of error in metadata assignation, motivated us to create a quality checking to determine the specie of an RNA-seq sample. In Rubinoff et al. (30), the usage of mitochondrial DNA for phylogenetic and taxonomic inference was discussed and 2 extreme viewpoints emerged: using exclusively the mitochondrial DNA or fully excluding it. It appears that mitochondrial DNA does not fully answer or impairs the perspectives of advanced phylogenetics. However, the “mitochondrial barcode” approach does show an interesting gene marker: the mitochondrially encoded cytochrome c oxidase I (MT-CO1) (31), that could be sufficient for a quick check of the specie of an RNA-seq data. Indeed, this gene is highly expressed and reference sequences from many distinct species of animals are available. Thus, we selected specific k-mers with Kmerator, at the gene level, for MT-CO1. We repeated the procedure for MT-CO1 orthologs in different species, principally found in SRA database, using the appropriate species reference genome and transcriptome. These k-mers have been then quantified in three public data by species to check the efficiency of their usage. As shown in the Figure 4c, the research of MT-CO1 k-mers alone can discriminate most of the common Ensembl species and can be usable for a quick quality-check. However, without proper experiments we cannot support its usage with phylogenetically close species.

To conclude, we developed KmerExploR to rapidly control RNA-seq raw data quality and filter samples on unusual profiles or presence of contaminations. KmerExploR is a tool that provides a modular set of analyses like fastQC (https://qubeshub.org/resources/fastqc). It can be used in a complementary way to fastQC analysis to complete missing metadata in public datasets or to give a quick profile of the RNA-seq contents. The modular analysis is based on a k-mer selection from predictor genes, included into KmerExploR. The tool can be used to control any human RNA-seq dataset, and it can also be easily modified adding any other modular fonction.

### KmerExplor, an advanced usage for the detection of genomic or transcriptomic events

The above “checking application” of KmerExploR demonstrated all its potential in the rapid exploration of large public RNA-seq datasets before performing any biological query. However, the KmerExploR tool can also be used in a more advanced way such as biomarker search or discovery in human health. This application is a powerful one as it can compensate for the lack of completeness in genomic or transcriptomic references and we currently know that much important information may be missed by ignoring the unreferenced RNAs diversity (32). As a proof of concept we used a set of k-mers designed with Kmerator to identify events outside reference annotations including fusion or chimeric RNA, oncogene mutations and new long non coding RNA expression. We then applied k-mers quantification in a tumoral and non tumoral data set to evaluate the specificity and performance of the approach. The results obtained with a part of the Leucegene cohort are presented in Figure 5.

**Figure 5.** KmerExploR advanced usage: quantification of transcriptomic events outside the annotations. All barplots are represented for Dataset-LEUCEGENE described in table S1 (131 paired-end samples). This dataset includes normal CD34 positive cells as control (in green on the X axis) and different AML subtypes (in black on the X axis). For each barplot, the legend lists the set of predictors for which k-mers mean counts normalized per billion (Y axis) are computed. (a) Chimeras detection. 2 well-known fusion RNAs associated with chromosomal translocation and their reciprocal counterparts are presented: RUNX1-RUNXT1 t(x,21) and PML-RARA t(15, 17). (b) Mutations detection. TET2, KRAS and CEBPA genes are used in AML diagnosis. The barplot shows 4 different mutations for these genes, detected specifically into some AML samples. The reference allele for each of these mutations is detected in almost all samples. (c) new lncRNA detection: NONE “chr2-p21” lncRNA described in Ruffle et al. This transcript is expressed in the whole dataset but shows different levels of expression depending on AML subtype.

The selection includes different AML subtypes and normal CD34 positive cells as control (Dataset-LEUCEGENE described in table S1). The results obtained with two well-known fusion RNAs associated to chromosomal translocation, RUNX1-RUNXT1 (t(x,21)) and PML-RARA (t(15, 17)) and their reciprocal counterparts RUNXT1-RUNX1 and RARA-PML are presented in Figure 5a. In this case, the k-mers, once designed by Kmerator, are restricted to those spanning the fusion jonction with at least 10 nucleotides in gene 1 or gene 2 of the fusion. All the normal CD34+ cells are negative and we only observe an expression in corresponding positive AML-subtypes. Figure 5b illustrates the results obtained for mutations in TET2, KRAS and CEBPA genes currently used in AML diagnosis. Once again, we only observe the presence of these mutations in positive samples, demonstrating the high specificity of the approach by k-mers. The expression of a new lncRNA was also quickly searched in the Leucegene dataset (see Figure 5c), we observe an homogeneous and low expression in CD34 normal cells compared to an heterogeneous one in AML subtypes. This lncRNA candidate was already described in Ruffle et al. (18), using for the first time the “k-mer concept” for checking new biomarker candidates, and we have demonstrated a restricted expression of the NONE “chr2-p21” lncRNA in the haematopoietic lineage using Leucegene and ENCODE datasets. Hence, for lncRNA candidates, following their discovery in a tissue/disease type, their specificity could be easily evaluated through quantification in a wide range of RNA-seq data including normal and pathological conditions as recently described by Riquier et al. (33).

In conclusion, the high specific expression of transcriptional events may lead them to be used as biomarkers for biological and health applications including cell therapy, diagnosis, prognosis or patient follow-up as it is already done with fusion RNAs and mutations.

## DISCUSSION

Considering the growing number of RNA-seq data, the use of raw data sequences is an important step to check with RNA-seq protocols or bioinformatic pipelines bias. Here, we demonstrated that the Kmerator Suite is an efficient and useful set of tools to verify RNA-seq quality and control intrinsic method and biological characteristics that often failed in technical description. We also showed that the Kmerator Suite can be used to quantify gene/transcript specific expression as well as to explore sequence variations at the transcriptional level. We first adapted the tool to human/mouse data ENSEMBL entry but it could be easily used with minor implementation to other species or references entries like GENCODE.

The meta-analyses performed in the present study with KmerExploR are a proof of concept of the procedure potential and could be extended to other biological RNA-seq questioning: i/ to extend the application to an enlarged set of microorganisms including new ones like SARS-Cov2 detection, ii/ to search for immuno-phenotyping profile in cancer datasets as already published by Mangul et al. (34). Considering advanced applications, we also demonstrated the potential of k-mers to explore gene expression in RNA-seq to reinforce biological questions or biomarker usage and discovery. Moreover, many other requests could be easily considered for annotated genes exploration like gene coexpression, or to compensate the lack of completeness in genomic or transcriptomic references to cover unreferenced RNA diversity and search for new spliced events, introns retention or new transcript categories including circular RNAs. In order to increase the potential of the k-mers approach, access to very large-scaling datasets like SRA level (> 40 000 samples) could be considered with efficient indexing structures development (35).

## DATA AVAILABILITY

RNA-seq libraries were downloaded from the European Nucleotide Archive of the European Bioinformatics Institute (ERR) (36). The reference GRCh38 genome and Ensembl 91 transcripts were downloaded from Ensembl. Kmerator is distributed under the MIT license. The Kmerator, KmerExploR and countTags softwares, documentation, and supplemental material presented herein are available from https://github.com/Transipedia/kmerator, https://github.com/Transipedia/kmerexplor and https://github.com/Transipedia/countTags respectively.

## SUPPLEMENTARY DATA

Supplementary table S1

## ACKNOWLEDGEMENTS

The authors are grateful to Henrietta Lacks, now deceased, and to her surviving family members for their contributions to biomedical research. The HeLa cell line that was established from her tumor cells without her knowledge or consent in 1951, have made significant contributions to scientific progress and advances in human health. The authors are also grateful to Rayan Chikhi for his comments and corrections.

## FUNDING

This work was supported by the Agence Nationale de la recherche for the project “Transipedia” [ANR-10-INBS-09]; the Canceropole Grand-Sud-Ouest “Trans-kmer” project [2017-EM24]; and the Region Occitanie for the project “SuriCare” [R19073FF].

### Author’s contribution

SR and TC designed the study. SR, CB and TC wrote the manuscript. ALB, NG and DG were contributors in the design of the study and manuscript corrections. SR developed the code of Kmerator, selected and downloaded public datasets, analysed data. BG and CB participated to Kmerator code improvements. CB analysed RNA-seq data and generated figures. BG developed KmerExploR code, generated k-mers countings and figures. JA and AB computed and corrected countTags. FR validated the RNA-seq data for mutation and chimeric RNAs, and helped for results interpretation. HX participated in Kmerator testing and checking. All authors read and approved the final manuscript.

