## Supplementary figures for "Kmerator Suite: design of specific k-mer signatures and automatic metadata discovery in large RNA-Seq datasets"

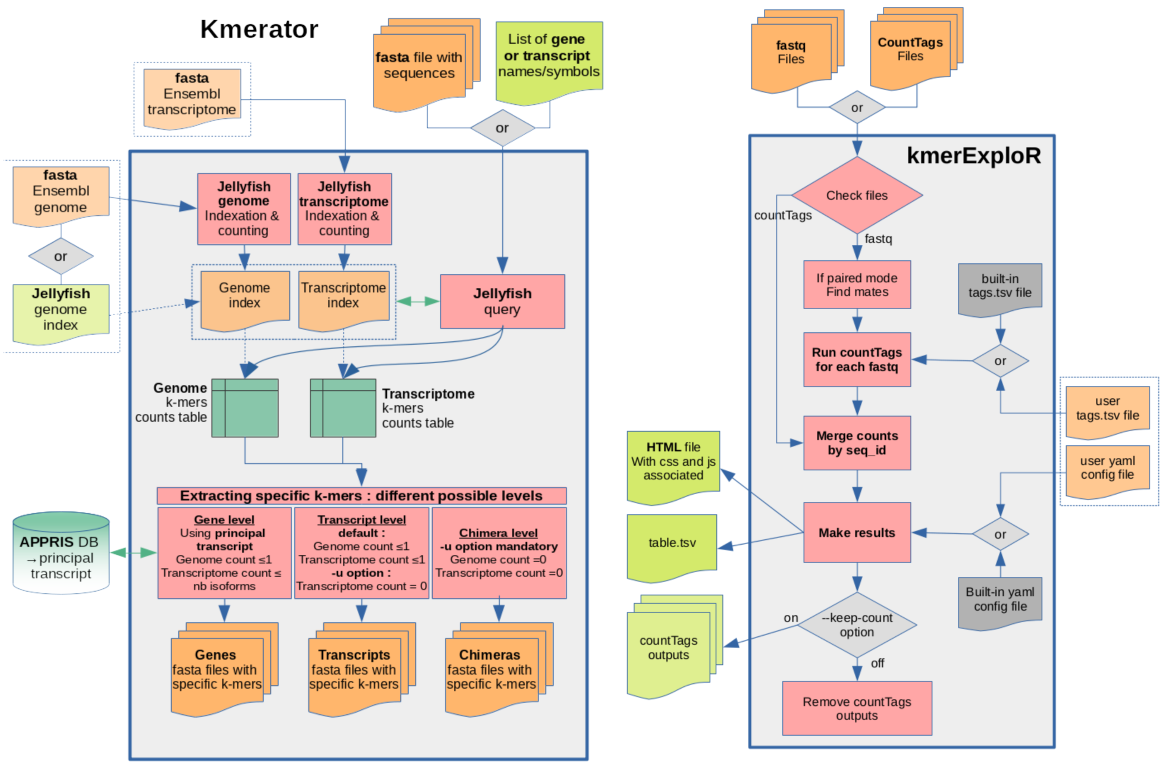


(a) (b)

Figure S1: **Kmerator and KmerExploR diagrams.** **(a)** Kmerator input is either a fasta file or a list of gene/transcript names. It also takes an Ensembl reference transcriptome (fasta format) and a reference genome (fasta or jellyfish format). In a first step, Kmerator uses jellyfish to index the transcriptome and, if necessary, the genome. In a second step and depending on the input, Kmerator extracts sequences corresponding to the given list (gene/transcript) or takes directly the sequences (fasta), and it generates k-mers count tables using jellyfish query in both the genome and the transcriptome. The third step is to extract the specific k-mers according to the chosen level (gene, transcript or chimera) and options (appris database, unnaotated). The output is a fasta file of specific k-mers for each input sequence/gene/transcript. **(b)** KmerExploR input is a set of fastq files (or a set of countTags output files). In the first instance, KmerExploR run countTags for each fastq, counting k-mers from the built-in tags file or from a specific user tags file. The k-mer counts are then averaged by id and merged in a single tsv file for all the samples. The outputs are an HTML file, a tsv file with the mean k-mer counts and countTags output if –keep-count option is on. The HTML page presents the graphical quantification results and strictly depends on the config file (built-in or user) where categories to show are defined.

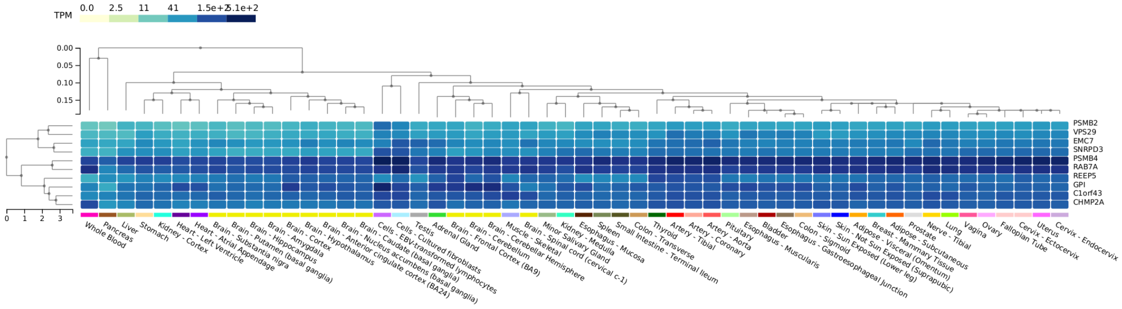


(a)

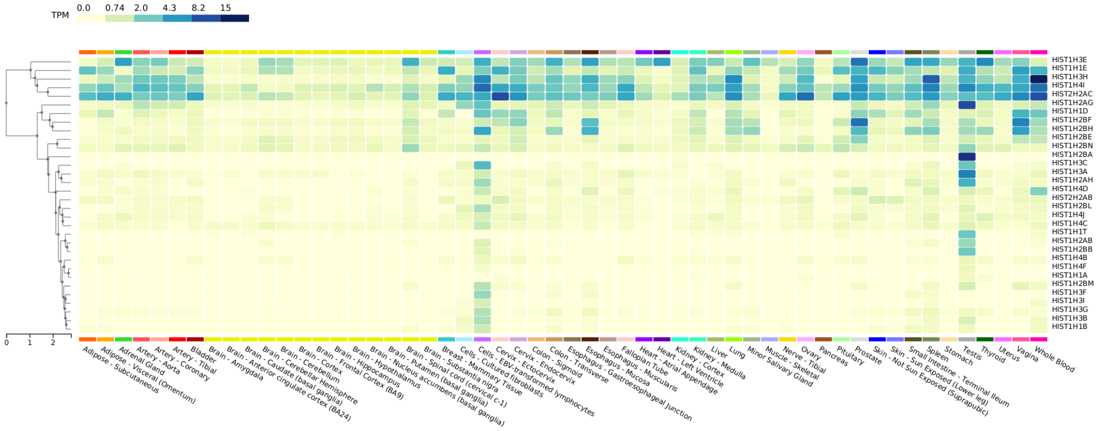


(b)

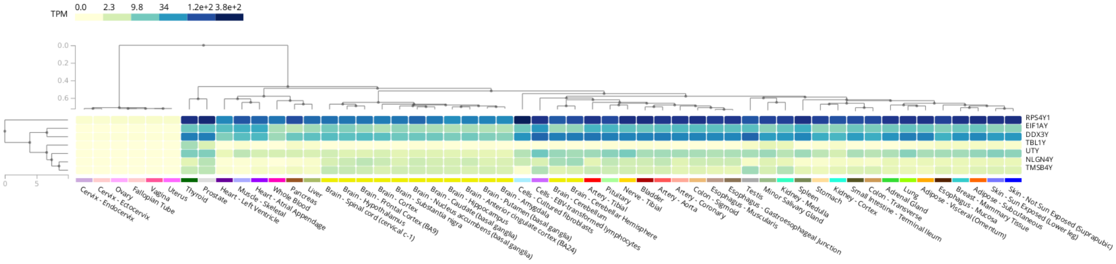


(c)

Figure S2: **GTEx expression heatmaps.** To help us define our predictor genes lists, we used the GTEx resource. Each heatmap downloaded from the GTEx portal represents the expression patterns across 54 different tissues (17382 samples, GTEx). The color scale is the expression in TPM. **(a)** Expression pattern heatmap for our selected ubiquitous genes. **(b)** Expression pattern heatmap for selected histone genes. **(c)** Expression pattern heatmap for Y chromosome specific genes.

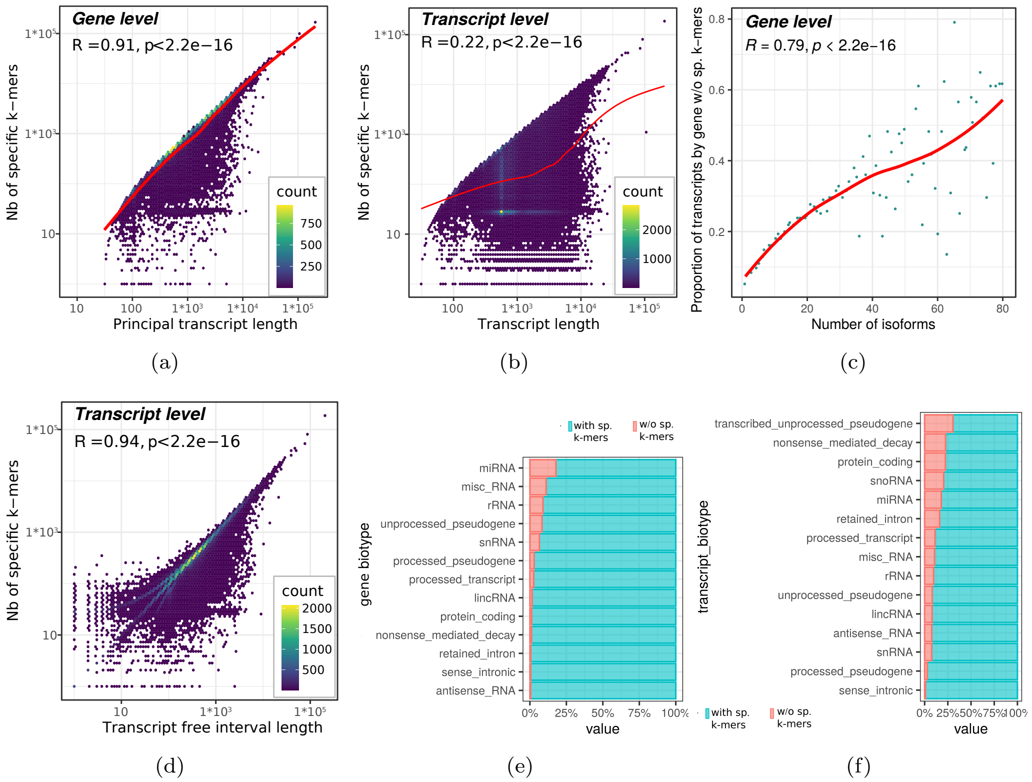


Figure S3: **Kmerator on the whole transcriptome. (a-b)** On the simulated data, we computed the Spearman correlation between the number of specific k-mers (Y axis) and the transcript length (X axis) at both gene (a) and transcript (b) levels. For the gene level, the length of the principal transcript (APPRIS database) is taken. Each point on the graph is a transcript and the color scale depends on the transcript density on the graph. **(c)** We also represented for each gene, the proportion of its associated transcripts without specific k-mers (Y axis), function of the number of isoforms and calculated the Spearman correlation. **(d)** We finally represented the number of specific k-mers (Y axis) function of the free interval length (X axis) at the transcript level. **(e)** For all the genes of the Ensembl annotation, we used Kmerator to compute specific k-mers (gene level) and we looked at the biotype of the genes having at least 1 specific k-mer (turquoise) versus the ones without specific k-mers (red). The small genes (miRNA, miscRNA) have the highest percentage without specific k-mers ( 20%). **(f)** In the same way, we computed specific k-mers for all the transcripts of the Ensembl transcriptome and looked at the biotype of the transcripts having at least 1 specific k-mer (turquoise) versus the ones without specific k-mers (red). Pseudogene transcripts, which are highly repetitive, are the ones having the highest proportion without specific k-mers ( 30%).

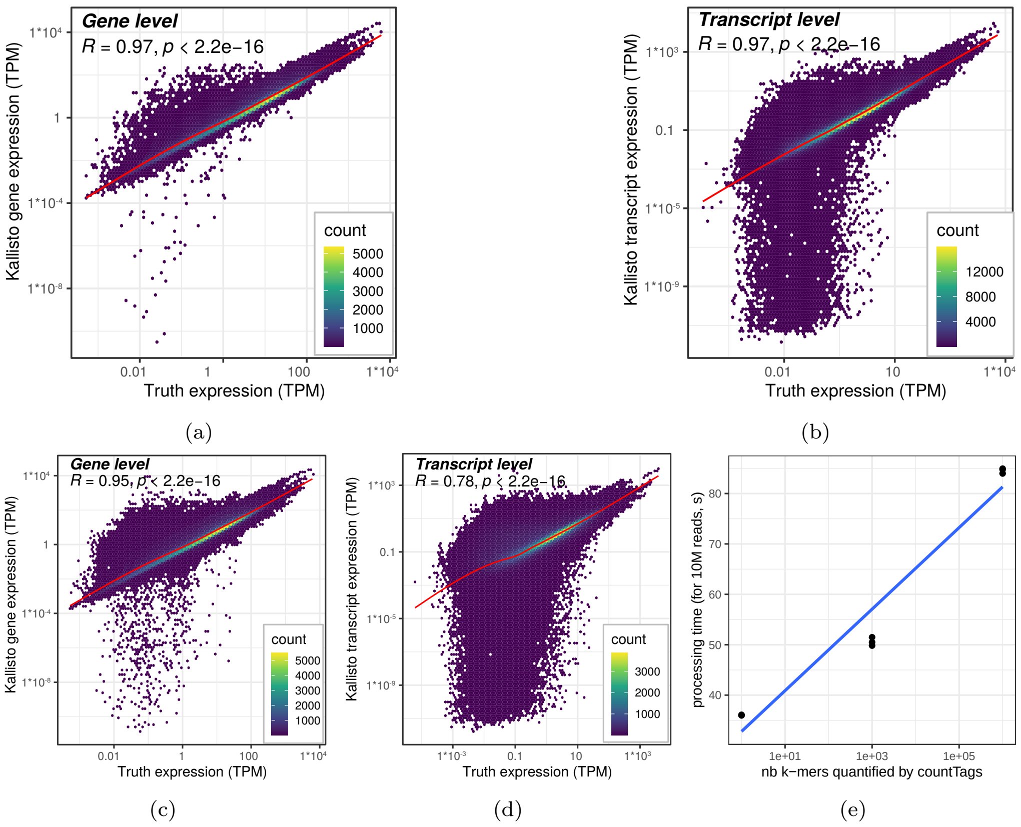


Figure S4: **Kallisto on the whole transcriptome. (a-b)** For the simulated data and the same gene/transcript set as represented with Kmerator (Figure 2), we computed Kallisto expression in TPM (Y axis) function of the true expression in TPM (X axis), for both genes (a) and transcripts (b). **(c-d)** For the remaining genes/transcripts not covered by Kmerator, we also computed Kallisto expression function of the true expression, again for both genes (c) and transcripts (d). Interestingly, the precision of Kallisto decreases on these gene/transcript sets. **(d)** We tested countTags processing time on a random selection of 10M reads of one simulated sample (Y axis). Processing time depends on the number of input k-mers (X axis) requested: 1, 1000 or 1M k-mers.

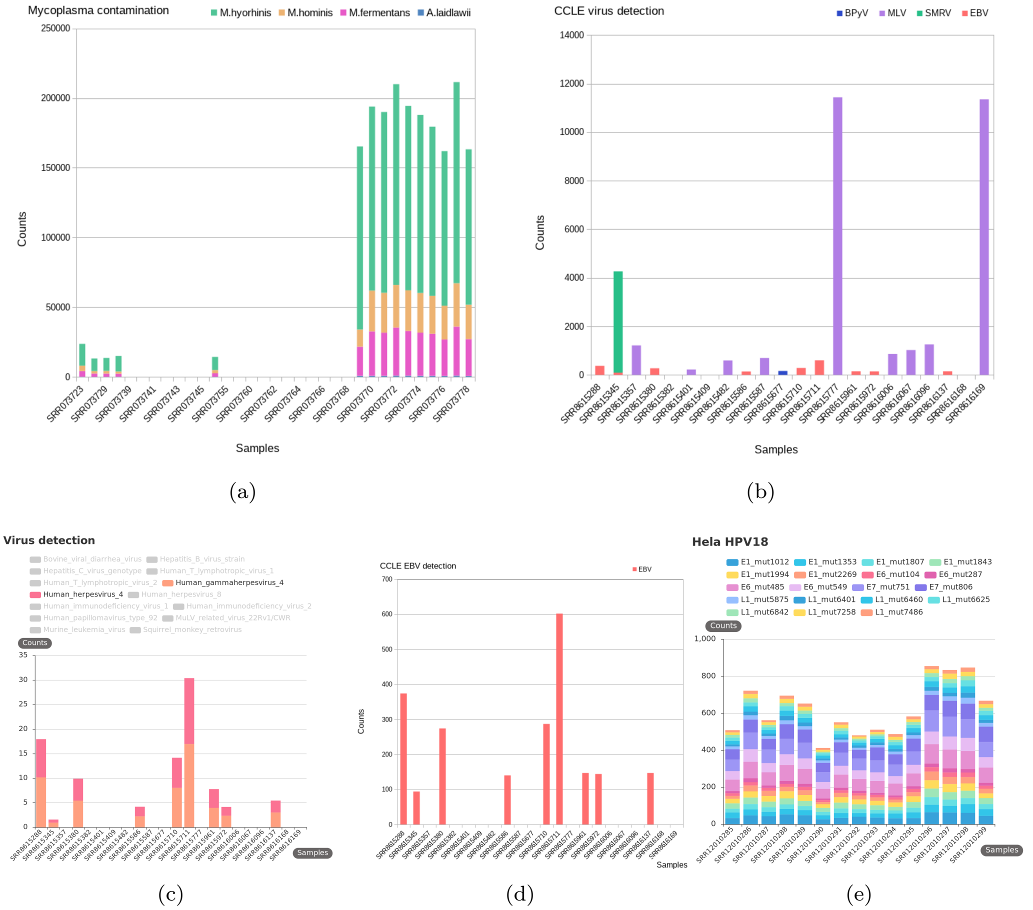


Figure S5: **Contaminations. (a)** Barplots generated from Olarerin-George et al. study, representing the number of reads/million (Y axis) mapping to 4 species of mycoplasma : A.laidlawii PG-8A, M.fermentans M64, M.hominis ATCC 23114 and M.hyorhinis MCLD, in a subset of the PRJNA153913 RNA-seq samples (Dataset-MYCO). **(b)** Barplot of the number of reads mapping to the viruses determined by Taxonomer (Y axis), from the Uphoff et al. study, on the samples from the Dataset-VIRUS-CCLE described in table S1 (X axis). **(c)** Barplot of the Epstein-Barr virus (EBV, gammaherpesvirus4) and herpesvirus4 contaminations determined with KmerExploR. The EBV is one of the most common human viruses. Samples are on the X axis and mean k-mer counts normalized per billion of k-mers on the Y axis. **(d)** In comparison, we generated the barplot of the number of reads (Y axis) mapping to the EBV from Uphoff et al. (2019). **(e)** HeLa determination on the Dataset-HELA-SUPP (15 paired-end samples) with KmerExploR. Samples are on the X axis and mean k-mer counts normalized per billion of k-mers on the Y axis.
